# Parameterising continuum models of heat transfer in heterogeneous living skin using experimental data

**DOI:** 10.1101/354563

**Authors:** Sean McInerney, Elliot J Carr, Matthew J Simpson

## Abstract

In this work we consider a recent experimental data set describing heat conduction in living porcine tissues. Understanding this novel data set is important because porcine skin is similar to human skin. Improving our understanding of heat conduction in living skin is relevant to understanding burn injuries, which are common, painful and can require prolonged and expensive treatment. A key feature of skin is that it is layered, with different thermal properties in different layers. Since the experimental data set involves heat conduction in thin living tissues of anesthetised animals, an important experimental constraint is that the temperature within the living tissue is measured at one spatial location within the layered structure. Our aim is to determine whether this data is sufficient to reliably infer the heat conduction parameters in layered skin, and we use a simplified two-layer mathematical model of heat conduction to mimic the generation of experimental data. Using synthetic data generated at one location in the two-layer mathematical model, we explore whether it is possible to infer values of the thermal diffusivity in both layers. After this initial exploration, we then examine how our ability to infer the thermal diffusivities changes when we vary the location at which the experimental data is recorded, as well as considering the situation where we are able to monitor the temperature at two locations within the layered structure. Overall, we find that our ability to parameterise a model of heterogeneous heat conduction with limited experimental data is very sensitive to the location where data is collected. Our modelling results provide guidance about optimal experimental design that could be used to guide future experimental studies.

**Nomenclature:** A brief description of all variables used in the document are given in Table 1.

Table 1:
Variable nomenclature and description.

Table 1: Variable nomenclature and description.
| Variable | Description |
| --- | --- |
| $t$ | time |
| $t_j$ | $j^{\text{th}}$ sample in the time series |
| $t_c$ | critical time |
| $\delta t$ | time between samples |
| $x$ | depth below the surface of skin |
| $l_1$ | depth of the interface between the skin and fat layers |
| $l_2$ | depth of the bottom of the fat tissue |
| $p$ | depth of first probe |
| $q$ | depth of second probe |
| $T$ | non-dimensional temperature |
| $T_1$ | non-dimensional temperature in skin layer |
| $T_2$ | non-dimensional temperature in fat layer |
| $\widehat{T}$ | non-dimensional synthetic temperature data, obtained using $(\widehat{D}_1, \widehat{D}_2)$ |
| $\mathcal{T}_1$ | dimensional temperature in skin layer |
| $\mathcal{T}_2$ | dimensional temperature in fat layer |
| $\mathcal{T}_0$ | dimensional initial temperature in the experiments |
| $\mathcal{T}_h$ | dimensional temperature in the scald creation device |
| $\widehat{\mathcal{T}}$ | dimensional experimental temperature data |
| $D_1$ | thermal diffusivity of skin |
| $D_2$ | thermal diffusivity of fat |
| $D_{\text{eff}}$ | homogenised averaged thermal diffusivity |
| $\widehat{D}_1$ | target thermal diffusivity of skin |
| $\widehat{D}_2$ | target thermal diffusivity of fat |
| $D_1^{(\min)}$ | minimum value of $D_1$ |
| $D_1^{(\max)}$ | maximum value of $D_1$ |
| $D_2^{(\min)}$ | minimum value of $D_2$ |
| $D_2^{(\max)}$ | maximum value of $D_2$ |
| $d$ | non-dimensional temperature discrepancy between model and data |
| $\varepsilon$ | discrepancy threshold |
| $\mathcal{I}_1$ | indicator function for one probe |
| $\mathcal{I}_2$ | indicator function for two probes |
| $A$ | area of the bounded $(D_1, D_2)$ parameter space |
| $\mathcal{A}_1$ | proportion of the bounded $(D_1, D_2)$ parameter space where $\mathcal{I}_1 = 1$ |
| $\mathcal{A}_2$ | proportion of the bounded $(D_1, D_2)$ parameter space where $\mathcal{I}_2 = 1$ |

## 1. Introduction

Injuries caused by accidental exposure to hot liquids are common, painful and often require extensive long-term treatment [1]. To improve our understanding of how thermal energy propagates through human skin, experimental studies often work with porcine (pig) skin because porcine skin is anatomically similar to human skin [2–8]. Many experimental studies deal with heat conduction in excised non-living tissues [6, 7, 9, 10]. In contrast, the experimental protocols developed by Cuttle and colleagues [11–14] are unique because they quantify heat conduction in living porcine tissues. Working with living tissues is far more biologically relevant than working with excised non-living tissues. Cuttle’s experimental protocol involves working with anesthetised living pigs that are given analgesia. A thermocouple probe, referred to as the *subdermal probe*, is inserted obliquely under the skin of the animal at various locations on the body [11–14]. To initiate an experiment, a cylindrical scald creation device is placed onto the surface of the skin so that the centre of the circular scald device is above the subdermal probe. Pre-heated water is pumped into the scald device and suctioned out of the device at an equal rate to ensure that a constant level of water at a particular temperature is maintained in the device at all times during the experiment. The temperature response in the living skin is measured by the subdermal probe as a function of time during the experiment. This time series data reveals information about how the thermal energy propagates through the living skin, and this experimental protocol can be used to study how thermal energy propagates through skin in different locations on the body. Further, by using pigs of different ages the same experimental protocol can be used to study how the propagation of thermal energy depends on skin thickness [2].

A visual summary of Cuttle’s experimental porcine model is given in Figure 1. The image in Figure 1(a) shows a portion of excised skin at the conclusion of an experiment highlighting the location and size of the subdermal probe. The histology image in Figure 1(b) highlights the layered structure of the skin. The epidermis and dermis forms the upper layer of the skin where hair follicles are present [15, 16]. The epidermis and dermis are bright pink in Figure 1(b), and throughout this study we treat the epidermis and dermis as a single layer that we call the *skin* layer. Underneath the skin layer there is a layer of fat that is a lighter shade of pink in Figure 1(b). Throughout this work we refer to this lower layer as the *fat* layer. As indicated in Figure 1(c), we adopt a coordinate system where *x* = 0 corresponds to the skin surface. The interface between the fat and skin is located at *x* = *l*_1_ > 0, we have *l*_1_ = 1.6 mm in this case. The interface between the fat and the underlying muscle and bone is at *x* = *l*_2_ > *l*_1_, and we have *l*_2_ = 4.0 mm in this case. Our conceptual idealisation of the structure of the layered tissues is given in Figure 1(c) where the subdermal probe is placed at *x* = *l*_2_ since experimental data reported by Cuttle involves placing the probe at the bottom of the fat layer [2, 17]. A summary of the kind of experimental data reported by Cuttle is given in Figure 1(d). In this particular experiment the probe is located at the interface of the fat and muscle, *x* = *l*_2_, and a scald creation device of diameter 50 mm is placed on the surface of the skin [2]. Water at temperature of 50°C is maintained in the scald creation device for a duration of 120 s, and the time series data showing the temperature at the subdermal probe is recorded, as shown. It is worth noting that the total depth of the tissue (4 mm) is much smaller than the diameter of the scald creation device (50 mm), so that 4/50 = 0.08 ≪1, as illustrated in Figure 1(e). Since the centre of the circular scald creation device is placed directly over the location of the probe the heat transfer downward through the skin can be idealised as a one-dimensional process [17].

**Figure 1:**
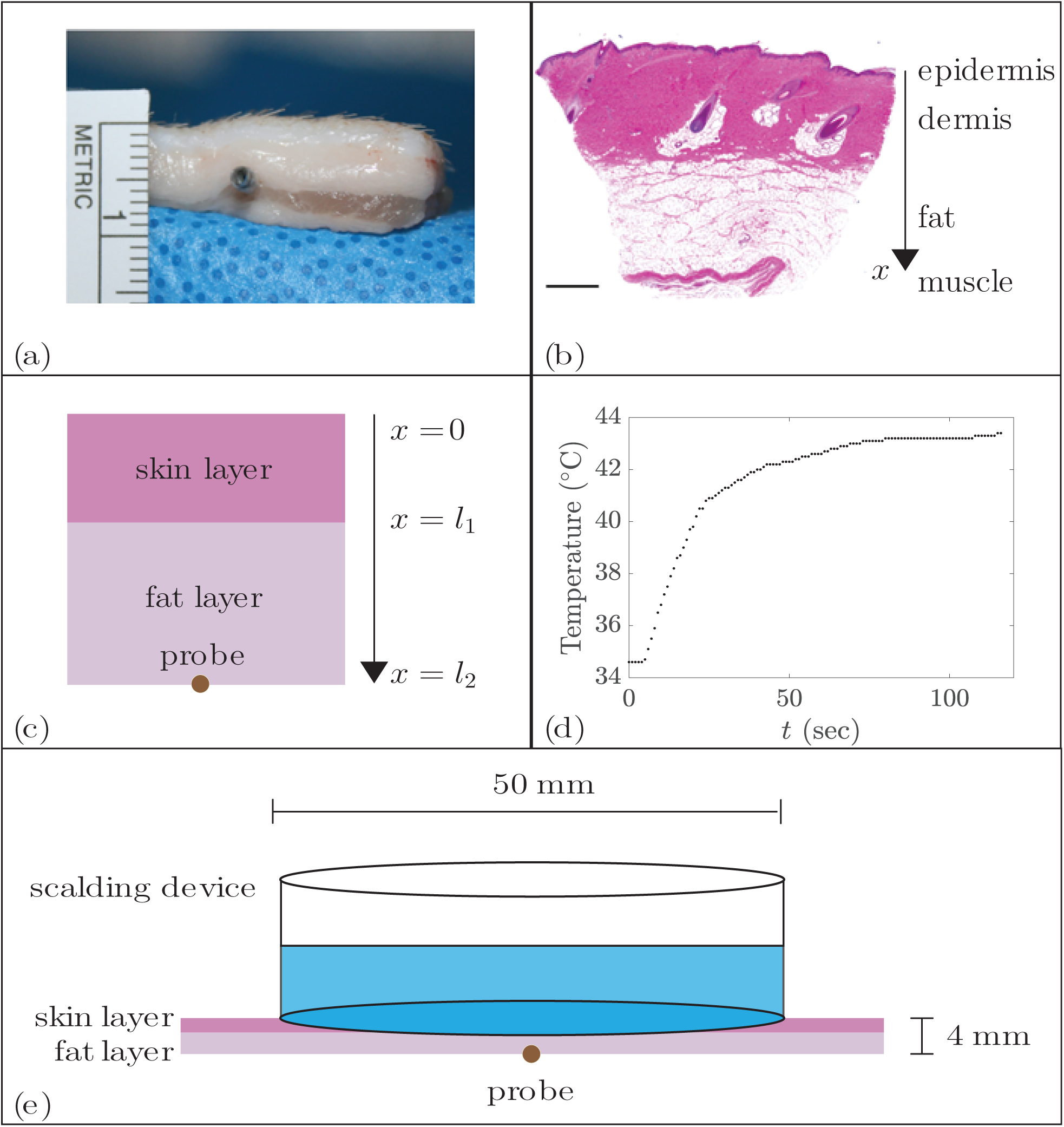
Developing a two-layer heterogeneous mathematical model to mimic the experimental porcine model. (**a**) Excised skin, showing the location of the probe and the depth of the tissue. (**b**) Histological image of normal porcine skin. Scale bar is 1 mm. The depth below the surface of the skin is denoted by *x* ≥ 0. (**c**) Conceptual two-layer model of the tissue with a skin layer (bright pink) sitting above the fat layer (lighter pink). The interface between the two layers is at *x* = *l*_1_, and the probe is located at the bottom of the fat layer, *x* = *l*_2_. (**d**) Example of the temporal variation of dimensional temperature, 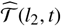, reported in [17]. Data is obtained from a subdermal temperature probe at *x* = *l*_2_. The water in the scald creation device is held at 50°C for a duration of 120 s. (**e**) Schematic showing that the tissues are very thin (4 mm) compared to the diameter of the scald creation device (50 mm). Images in (a) and (b) are reproduced from [17] with permission.

A prominent feature of the skin, highlighted in Figure 1(b), is the layered structure where we see that the fat layer is below the skin layer. This kind of histological information has been previously incorporated into mathematical descriptions of heat transfer in skin by explicitly accounting for the layered, heterogeneous structure of the tissue. These previous models have often been based on generalisations of Pennes’ bioheat equation [18–20] and re-formulated as a multilayer heterogeneous heat transfer model where the thermal properties can vary between the different layers [21–25]. A key limitation of working with such a heterogeneous multilayer heat transfer model is that they are more challenging to parameterise than simpler single layer models. This is a consequence of the fact that there are a greater number of unknown parameter values in a multilayer heterogeneous model compared to a simpler single layer model of heat transfer. This challenge is particularly acute if we consider parameterising a mathematical model of heat transfer using Cuttle’s realistic experiments that report the temperature response at one location within the layered structure. This experimental limitation is difficult to overcome because inserting multiple probes simultaneously at different depths would risk compromising the integrity of the living tissues. Our previous work has involved calibrating the solution of much simpler single layer homogeneous models to match data from Cuttle’s experiments [2, 17]. However, these previous studies suffer from the limitation that they implicitly treat the thermal parameters of the skin layer and fat layer together into a simplified, vertically averaged, homogenised single layer [26]. While this approach is mathematically convenient, it is unclear whether a single layer model is appropriate since we know that one of the main biological roles of the fat layer is to provide thermal insulation [27]. Therefore, we expect that the thermal properties of the skin and fat layers could be very different.

In this work we use a two-layer heterogeneous model to describe heat conduction in living tissues. Our aim is to perform a suite of synthetic experiments with realistic parameter values to mimic data generated by Cuttle’s experimental protocol. With this synthetic data we explore the extent to which we can confidently estimate the thermal diffusivity in each layer when we have limited experimental observations where the temperature is reported at one single location within the layered tissues. To achieve this, we use the solution of the two-layer heterogeneous model, parameterised with biologically-relevant estimates of the thermal diffusivity of skin and fat, to generate synthetic data that mimics Cuttle’s experimental protocol where a single probe is placed at the bottom of the fat layer. Given that the synthetic data is generated with known estimates of the thermal diffusivity in the skin and fat layers, we then systematically explore the parameter space to investigate whether the kind of data can be used to reliably determine parameters in the heterogeneous mathematical model. Once we have demonstrated how delicate this parameter estimation task can be, we turn our attention to the question of experimental design. First, we explore whether our ability to determine the parameters in the two-layer model varies when we alter the location of the single subdermal probe. Second, we explore the extent to which our ability to estimate the parameters improves when we consider synthetic experiments with two probes so that the temperature is recorded simultaneously at two different positions within the layered skin.

## 2. Mathematical model

We model the transfer of heat through the skin and fat layers using a one-dimensional model. This is a reasonable assumption given that the depth of the tissue is much smaller than the width of the scald creation device used in Cuttle’s experimental protocol [2, 17]. If the tissue depth was comparable to the diameter of the scale creation device it would be more appropriate to use a two- or three-dimensional mathematical model. In this work we assume that the temporal and spatial distributions of dimensional temperature in the skin layer, 𝒯_1_(*x*, *t*), and the fat layer 𝒯_2_(*x*, *t*), are governed by

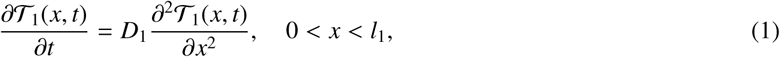

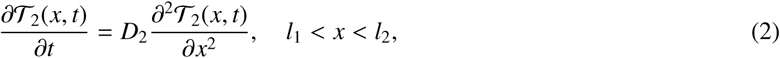

where *D*_1_ > 0 is the thermal diffusivity of the skin and *D*_2_ > 0 is the thermal diffusivity of fat. We have not included any source terms in Equations (1)-(2). Although some previous studies have incorporated source terms to account for the transfer of thermal energy from the skin tissues to the the blood supply [18], known as perfusion, our previous work, in which we calibrated the solution of a single layer model to match data from Cuttle’s experiments suggests that the role of perfusion is negligible in these experiments [17]. We note that the assumption that perfusion plays a negligible role has also been adopted in other modelling studies [21].

Experimental data suggests that the initial variation in temperature with depth is negligible [2]. Therefore we choose the initial condition to be

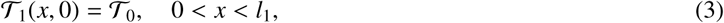

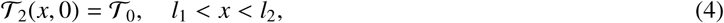

where 𝒯_0_ is the initial dimensional temperature of the skin and fat layers.

The boundary condition at *x* = 0 corresponds to the placement of the scald creation device on the skin surface. Cuttle’s experimental protocol carefully maintains a constant temperature in the scald creation device by pumping water of a constant temperature into the device at the same rate as water is pumped from the device, thus ensuring the maintenance of a constant temperature at the skin surface [2]. Therefore, we represent this as a Dirichlet boundary condition at *x* = 0. For simplicity, we assume that the flux of thermal energy at the base of the fat layer, *x* = *l*_2_, is negligible and we will comment on the validity of this assumption in Section 4. Together, these boundary conditions are incorporated into the model by specifying

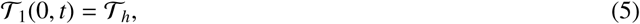

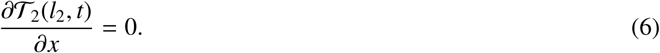

where 𝒯_*h*_ is the dimensional temperature of the water in the scald creation device.

In the literature, there are several different interface conditions that can be implemented in multilayer models of heat transfer [28–32]. Here we take the simplest, most fundamental approach by assuming perfect contact between the skin and fat layers. This amounts to assuming that the temperature and the flux of thermal energy are continuous at the interface

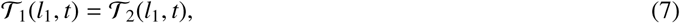

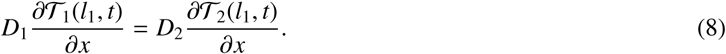

An attractive feature of Cuttle’s experimental design is that the temperature of the water in the scald creation device can be easily altered [2, 17]. For example, this experimental protocol has been used previously to study how skin responds to different temperature burns by using water at 50°C, 55°C and 60°C in the scale creation device [2]. Therefore, to ensure that our analysis can easily incorporate this feature of the experiments we non-dimensionalise the dependent variable in Equations (1)-(8) so that all of these different experimental conditions can be represented by the same mathematical model without explicitly considering the role of 𝒯_*h*_. To non-dimensionalise the dependent variable we introduce

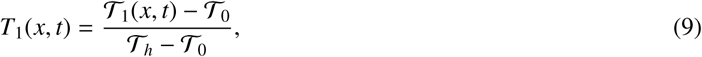

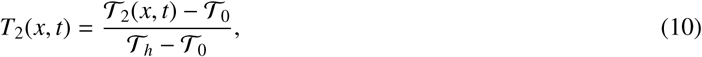

where *T*_1_(*x*, *t*) ∈ [0, 1] is the non-dimensional temperature in the skin layer and *T*_2_(*x*, *t*) ∈ [0, 1] is the non-dimensional temperature in the fat layer. Re-writing the governing equations in terms of these non-dimensional variables gives

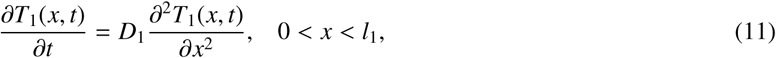

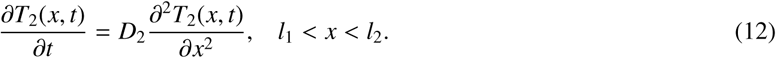

The initial condition for the non-dimensional model is

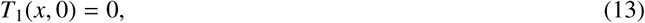

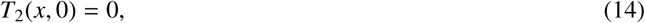

and the relevant boundary conditions are

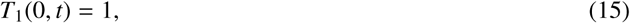

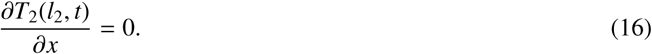

Finally, the interface conditions are written as

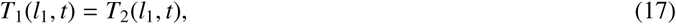

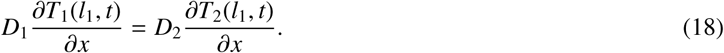

Equations (11)-(18) constitute the mathematical model that we consider in this study. For any particular choice of *D*_1_ and *D*_2_, the model can be solved to predict the temporal and spatial distribution of non-dimensional temperature within the two-layer problem, *T*_1_(*x*, *t*) and *T*_2_(*x*, *t*). These non-dimensional temperature profiles can be re-scaled, according to Equations (9)-(10), to give 𝒯_1_(*x*, *t*) and 𝒯_2_(*x*, *t*), which represent any particular experimental condition characterised by different choices of 𝒯_0_ and 𝒯_*h*_. A convenient feature of the mathematical model is that Equations (11)-(18) can be solved, very efficiently, using Laplace transforms [33]. This Laplace transform solution can be evaluated at little computational cost, regardless of the choice of *D*_1_, *D*_2_, *l*_1_ and *l*_2_. A full description of the Laplace transform solution technique, and validation of the accuracy of this approach is given in the Supplementary Material document. Algorithms and code used in this work are available at GitHub.

## 3. Results and discussion

Throughout this work we consider a fixed tissue geometry by setting *l*_1_ = 1.6 mm and *l*_2_ = 4.0 mm, which match the histology measurements in Figure 1(a)-(b). To solve Equations (11)-(18) we must specify *D*_1_ and *D*_2_. Our approach is to:

1. Select biologically-relevant estimates of the target parameters,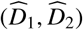;

2. Solve Equations (11)-(18) with the target parameters;

3. Extract time series data from the solution generated in Step 2 so that the synthetic data from the mathematical model is consistent with Cuttle’s experimental data;

4. Explore the solutions of Equations (11)-(18) across the (*D*_1_, *D*_2_) parameter space to assess how well we could estimate 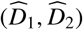 using the synthetic data generated in Step 3; and

5. Use the mathematical modelling tools to explore whether we can optimise the experimental design to improve our ability to reliably estimate 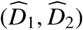.

### 3.1. Parameter inference: single probe

To generate synthetic data we must first estimate target parameters, 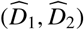. Previous work that interprets data from Cuttle’s experiments with a simplified, single-layer, homogenised mathematical model leads to an estimate of the homogenised effective thermal diffusivity, *D*_eff_ = 0.014 mm^2^/s. We will use this estimate to guide our choice of 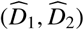. To achieve this we assume that *D*_eff_ corresponds to the homogenised thermal diffusivity of the two-layer tissue that is given by a weighted harmonic mean, 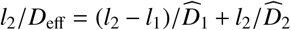. In addition, we make use of the fact that a key physiological role of the fat layer is to provide thermal insulation [27]. Therefore, we incorporate this into our heterogeneous multilayer model by requiring that *D*_2_ < *D*_1_, and we set 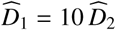 to reflect this. Combining these two biologically-motivated assumptions gives *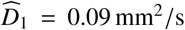* and 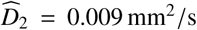, and we hold these target parameters constant throughout this work.

The solution of Equations (11)-(18) parameterised with 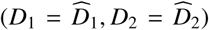 is shown in Figure 2(a). Here, the initial temperature is zero, and we see that energy is introduced into the system through the Dirichlet boundary at *x* = 0 mm, for *t* > 0. As the solution evolves, the temperature is continuous at the interface but the spatial gradient of temperature is discontinuous at the interface. From a modelling perspective, it is natural for us to visualise the entire spatial and temporal features of the solution of Equations (11)-(18) in Figure 2(a). However, this level of detail is not available in Cuttle’s experimental protocol [11–14] because temperature is measured at one spatial location only. Therefore, to ensure that the synthetic data we extract from the solution of Equations (11)-(18) is compatible with Cuttle’s experimental data, we use the solution of the mathematical model to generate time series data, showing *T*_2_(*l*_2_, *t*), at one spatial location only. At first we focus on *x* = *l*_2_ as the probe is placed at the bottom of the fat layer in the experiments [2, 17]. Later, in Section 3.2, we also consider the influence of varying the location of the probe.

**Figure 2:**
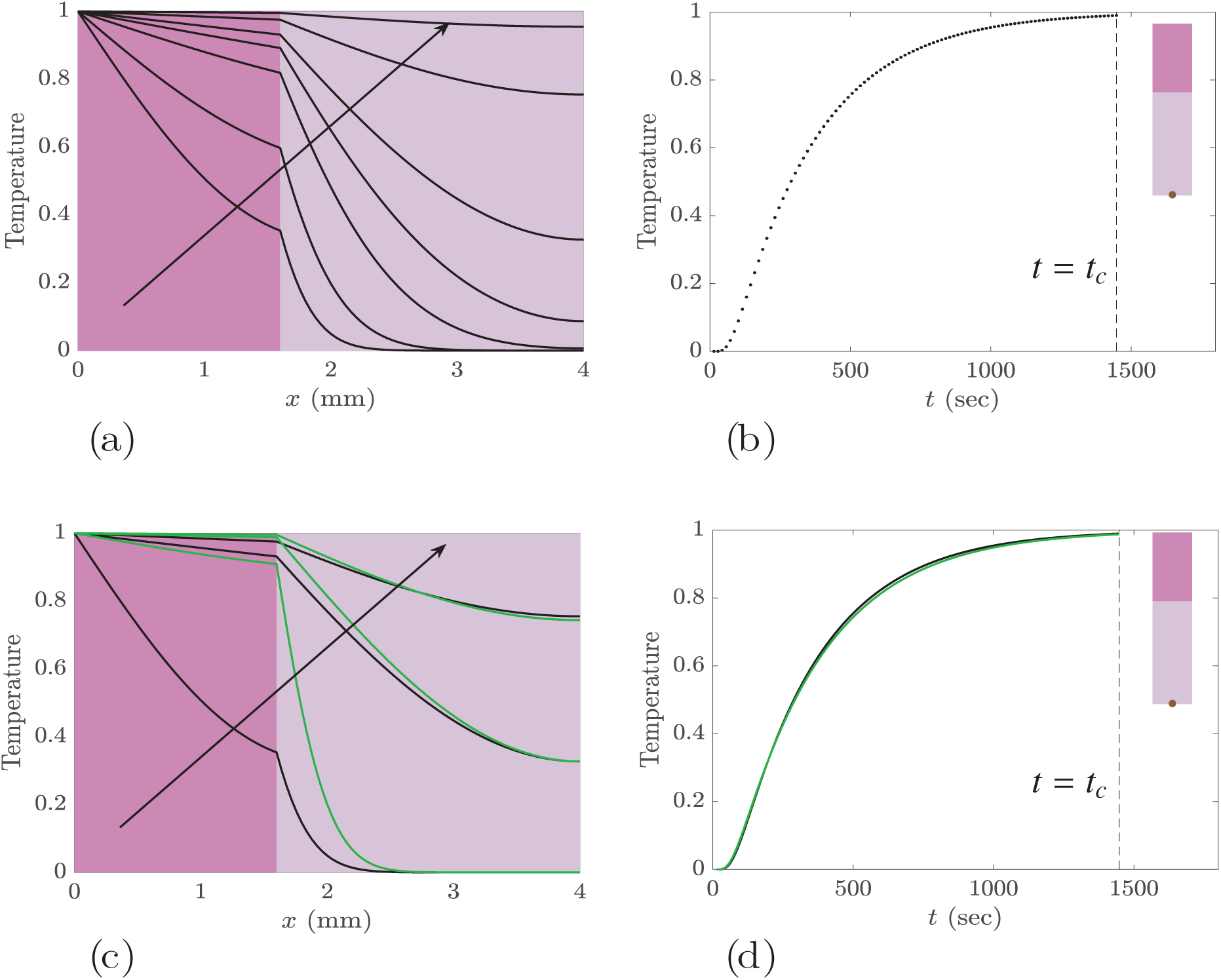
Solutions to Equations (11)-(18) for *l*_1_ = 1.6 mm and *l*_2_ = 4 mm. In (**a**) we set the thermal diffusivities to be the target parameters, *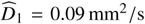* and 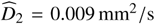. The solutions of Equations (11)-(18) are plotted at *t* = 10, 20, 50, 100, 200, 500 and 1000 s, with the arrow showing the direction of increasing *t*. (**b**) Synthetic time series data, 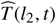, shows the temperature at the location of the probe, *x* = *l*_2_. The time series is constructed using 100 equally-spaced time point between *t* = *t*_*c*_/100 and *t* = *t*_*c*_, where *t*_*c*_ = 1447.5 s. (**c**) Comparison of *T*_1_(*x*, *t*) and *T*_2_(*x*, *t*) for two different parameter pairs. The black curves show the solution using the target parameters, *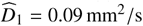* and *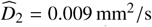*, and the green curves show solutions of the same model for a very different choice of parameters, *D*_1_ = 0.45 mm^2^/s and *D*_2_ = 0.0077 mm^2^/s. In (c) solutions are shown at *t* = 10, 200 and 500 s with the arrow showing the direction of increasing *t*. (**d**) Comparison of time series *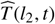* (black) and *T* (*l*_2_, *t*) (green), over the interval 0 ≤*t* ≤ *t*_*c*_, using the spatiotemporal solutions in (c). The bright pink and lighter pink background colours in (**a**) and (**c**) are chosen to correspond with the colour of the skin and fat layers in Figure 1(b). Subfigures (**b**) and (**d**) contain an inset showing the geometry of the skin layers with the brown circle showing the location of the probe, *x* = *l*_2_. In (**b**) and (**d**), the vertical dashed black line indicates the critical time, *t*_*c*_.

To construct the time series data from the solution of Equations (11)-(18), we must first decide on the interval of time that we will focus on. Since our aim is to estimate *D*_1_ and *D*_2_, it is useful to recall that an estimate of the duration of time required for the solution of Equations (11)-(18) to asymptote to the corresponding steady state solution will depend upon *D*_1_ and *D*_2_ [34–36]. This duration of time, called the *critical time* [37, 38], can be estimated by calculating the time required for the transient solution to reach within some small tolerance of the corresponding steady solution. For our choice of boundary conditions the long-time steady state solution of Equations (11)-(18) is 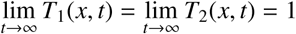. In this work we denote the critical time as *t*_*c*_, and we estimate the critical time by calculating *t*_*c*_ that satisfies *T*_2_(*l*_2_, *t*_*c*_) = 0.99, corresponding to a tolerance of 1%. For our values of *l*_1_, *l*_2_, *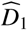* and *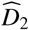* we have *t*_*c*_ = 1447.5 s. In this work we treat *l*_1_, *l*_2_, *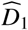* and *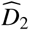* as constants which means that *t*_*c*_ is also a constant throughout this study. With our estimate of the critical time we generate the time series *T*_2_(*l*_2_, *t* _*j*_) with *t* _*j*_ = *jdt*, where *j* = 1, 2,…, 100 and *δt* = *t*_*c*_/100 s. This time series simply corresponds to 100 equally-spaced time points between *t* = *t*_*c*_/100 and *t* = *t*_*c*_, and we visualise this time series in Figure 2(b) for the problem shown previously in Figure 2(a). This time series confirms that *T*_2_(*l*_2_, 0) = 0, and *T*_2_(*l*_2_, *t*) approaches unity as *t* increases.

Now that we have specified the target parameters, 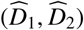, and defined how we extract synthetic data from the solution Equations (11)-(18), we explore how well we can estimate (*D*_1_, *D*_2_) so that the time series of *T*_2_(*l*_2_, *t*) matches the synthetic data. Ideally, there would be a unique choice of (*D*_1_, *D*_2_) for which the solution of the model matches the synthetic time series data. However, in practice we find there is large range of parameter pairs, (*D*_1_, *D*_2_), for which the time series data matches the synthetic time series data remarkably well. To illustrate this we show solutions of Equations (11)-(18) with very different choice of (*D*_1_, *D*_2_) in Figure 2(c). Here, we show the full spatial profile of the two solutions and it is obvious, from visual inspection alone, that the two solutions are very different. However, for these same two solutions, we see almost no difference when we view the time series, *T*_2_(*l*_2_, *t*), in Figure 2(d). This observation suggests that data provided by Cuttle’s experimental protocol might not be appropriate to constrain estimates of (*D*_1_, *D*_2_). This would be particularly challenging since Cuttle’s data will also be subject to experimental, biological and measurement variability that we have not accounted for in Figure 2. For simplicity and clarity, throughout this study we neglect the influence of such experimental variability, and we will comment on this assumption later, in Section 4.

Since the two time series in Figure 2(d) are difficult to visually distinguish, we introduce a discrepancy measure to assist in distinguishing between these time series quantitatively. In this work we use

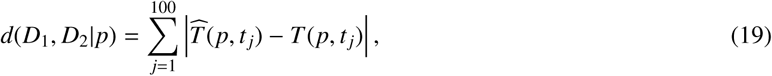

Where 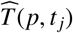 is the solution of Equations (11)-(18) parameterised with the target parameters 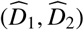, at location *x* = *p* and at time *t* = *t* _*j*_, and *T* (*p*, *t* _*j*_) is the solution of Equations (11)-(18), with some other choice of (*D*_1_, *D*_2_), at location *x* = *p* and at time *t* = *t* _*j*_. Here the sum is taken over 100 equally-spaced time points from *t* = *t*_*c*_/100 to *t* = *t*_*c*_ where *t*_*c*_ is first calculated for each choice of (*D*_1_, *D*_2_) that we consider. The key feature of this discrepancy measure is that there is a single probe at location *x* = *p*. Intuitively, we expect that choices of (*D*_1_, *D*_2_) that give rise to smaller values of *d*(*D*_1_, *D*_2_|*p*) could be reasonable estimates of 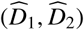. To help visualise which (*D*_1_, *D*_2_) parameter pairs lead to a close match with the synthetic data we use an indicator function

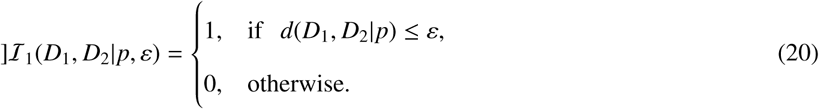

The indicator function is unity if the discrepancy is smaller than some specified threshold, *ε*, and zero elsewhere. Plotting *ℐ*_1_(*D*_1_, *D*_2_|*p*, *ε*) as a function of (*D*_1_, *D*_2_) is a useful way to visualise which combinations of (*D*_1_, *D*_2_) lead to good matches between *T* (*p*, *t*) and 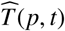. Plots of *ℐ*_1_(*D*_1_, *D*_2_|*p*, *ε*) as a function of (*D*_1_, *D*_2_) are constructed by discretising the (*D*_1_, *D*_2_) parameter space using a fine square mesh, and evaluating *ℐ* _1_(*D*_1_, *D*_2_|*p*, *ε*) at each point on the mesh for various choices of *ε*. All results presented in this work use a fine mesh of 2001 × 2001 equally-spaced values of *D*_1_ and *D*_2_. Each time we sweep across the parameter space we evaluate the solutions of Equations (11)-(18) more than 4 million times. Therefore, it is vitally important that the method we use to solve the governing equations is both accurate and efficient.

Results in Figure 3 show the region of (*D*_1_, *D*_2_) parameter space where *ℐ*_1_(*D*_1_, *D*_2_|*l*_2_, *ε*) = 1 for *ε* = 0.5, 1.0 and 1.5. Regardless of our choice of *ε*, we see that there are multiple combinations of (*D*_1_, *D*_2_) for which the time series, *T* (*p*, *t*), is very close to the synthetic time series, *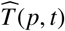*. As *ε* decreases, the area for which *ℐ*_1_(*D*_1_, *D*_2_|*l*_2_, *ε*) = 1 decreases, as the discrepancy measure is more restrictive. However, even with further reductions in *ε* > 0, we still observe a very large number of (*D*_1_, *D*_2_) pairs at which it is very difficult, if not impossible, to reliably distinguish between *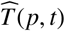* and *T* (*p*, *t*). This result indicates that relying solely upon Cuttle’s experimental protocol may not allow us to reliably identify unique choices of (*D*_1_, *D*_2_). We note that reducing the tolerance to zero, *ε* = 0, means that the region in (*D*_1_, *D*_2_) parameter space where *ℐ*_1_(*D*_1_, *D*_2_|*l*_2_, *ε*) = 1 does shrink to a unique point. However, working with a zero tolerance is impractical as we wish to allow for a small positive tolerance to account for some variability in the experimental measurements. Furthermore, working with a zero tolerance on a discretised parameter space is impractical since any regular meshing of the (*D*_1_, *D*_2_) parameter space would not precisely coincide with 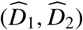.

**Figure 3:**
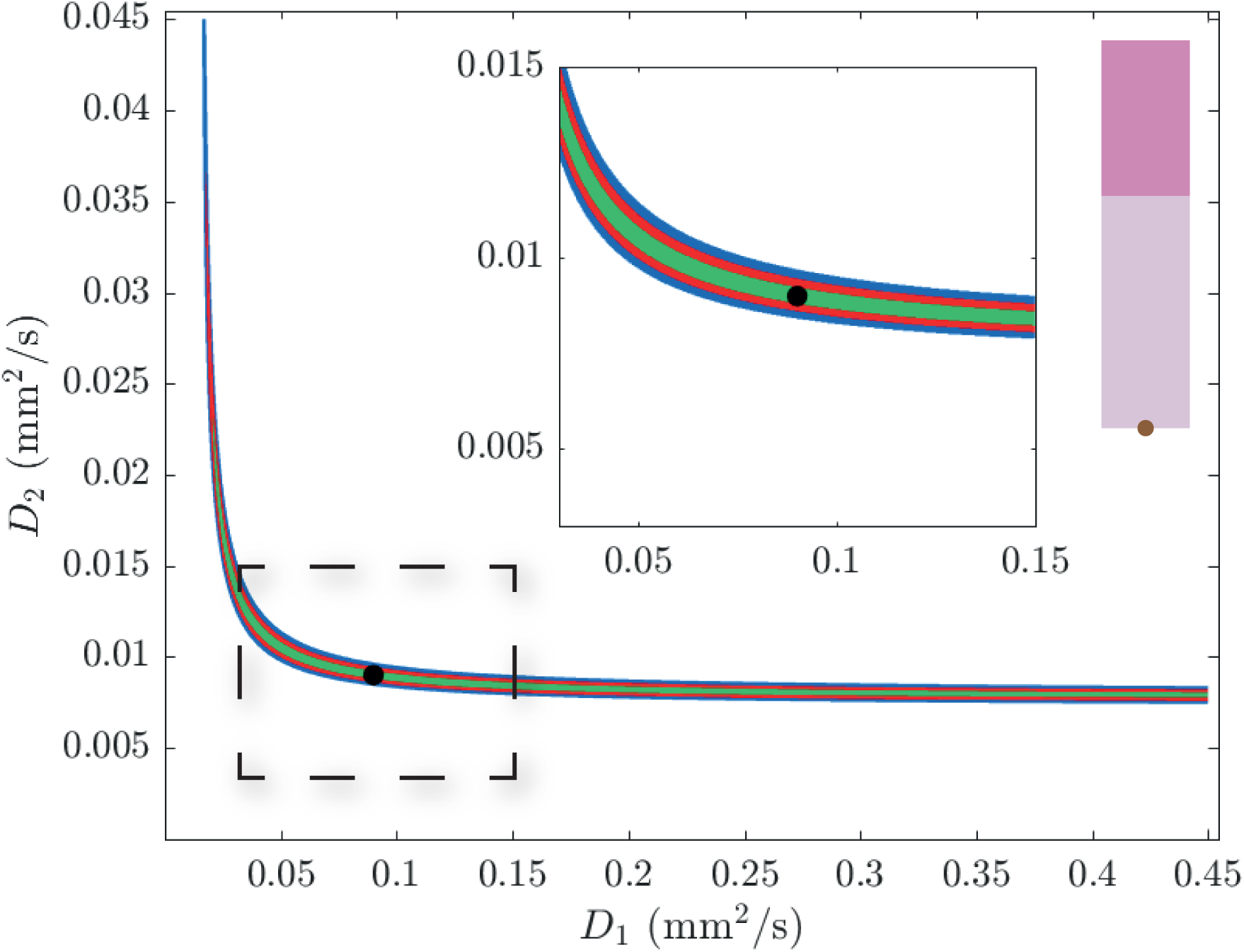
Regions of bounded parameter space where *ℐ*_1_(*D*_1_, *D*_2_|*l*_2_, *ε*) = 1, for *l*_1_ = 1.6 mm and *l*_2_ = 4 mm. The black circle indicates the target parameters, 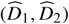. We plot *ℐ*_1_(*D*_1_, *D*_2_|*l*_2_, *ε*) on the bounded region 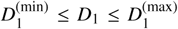 and *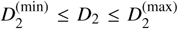*, where 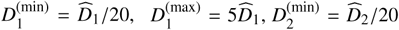 and 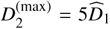. The coloured regions satisfy *ℐ*_1_(*D*_1_, *D*_2_|*l*_2_, *ε*) = 1 for *ε* = 0.5 (green), 1.0 (red) and 1.5 (blue). The central inset shows a magnified region, identified by the dashed rectangle in the main Figure, about the target parameter pair. The right-most inset indicates the geometry of the skin layers with the brown circle showing the location of the probe, *p* = *l*_2_.

While the plot of *ℐ*_1_(*D*_1_, *D*_2_|*l*_2_, *ε*) in Figure 3 is shown on the bounded region, *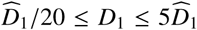* and, 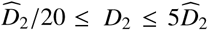 we also generated additional results by plotting *ℐ*_1_(*D*_1_, *D*_2_|*l*_2_, *ε*) over a larger support. These additional results (not shown) indicate that increasing the support leads to further choices of (*D*_1_, *D*_2_) pairs for which *T* (*p*, *t*) is very difficult to distinguish from *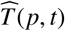*. That is, the extent of the coloured regions in Figure 3 continue to expand as the (*D*_1_, *D*_2_) support increases. This observation further corroborates our notion that it can be very difficult to infer (*D*_1_, *D*_2_), using a single probe located at *p* = *l*_2_ [17], and this observation motivates us to consider whether different choices of *p* could alter our ability to estimate (*D*_1_, *D*_2_).

### 3.2. Parameter inference: optimal placement of single probe

All results in Section 3.1 follow Cuttle’s experimental protocol by considering a single probe placed at *p* = *l*_2_. Therefore, we now repeat the process of generating the data in the same format as Figure 3 but for difference choices of probe location, *p*. To first explore the role of *p* we assume that some reasonable alternative choices to place the probe are:

1. the centre of the skin layer, *p* = *l*_1_/2,

2. the layer interface, *p* = *l*_1_;

3. the centre of the two-layer system, *p* = *l*_2_/2; and

4. the centre of the fat layer, *p* = *l*_1_ + (*l*_2_ − *l*_1_)/2.

Results in Figure 4 show plots that are equivalent to Figure 3 except that we consider these four different choices of *p*. Comparing data in Figures 3-4 shows that the choice of *p* has a dramatic impact upon the sensitivity of our ability to distinguish between *T* (*p*, *t*) and 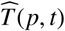. Perhaps the most obvious result is that choosing *p* = *l*_1_/2, as in Figure 4(a), leads to a very poor ability to estimate (*D*_1_, *D*_2_) since the extent of the coloured region is very large. This result makes intuitive sense because placing a single probe in the skin layer provides very little direct information about *D*_2_. In contrast, placing the probe at the centre of the two-layer system, *p* = *l*_1_ + (*l*_2_ − *l*_1_)/2, as in Figure 4(c), provides a better opportunity to estimate (*D*_1_, *D*_2_) since the extent of the coloured regions are smallest compared to the other choices of *p* in Figures 3-4. Overall, it appears to be optimal to place the probe in the fat layer, rather than the skin layer. This is a useful outcome as it is consistent with Cuttle’s experimental protocol [11–14].

**Figure 4:**
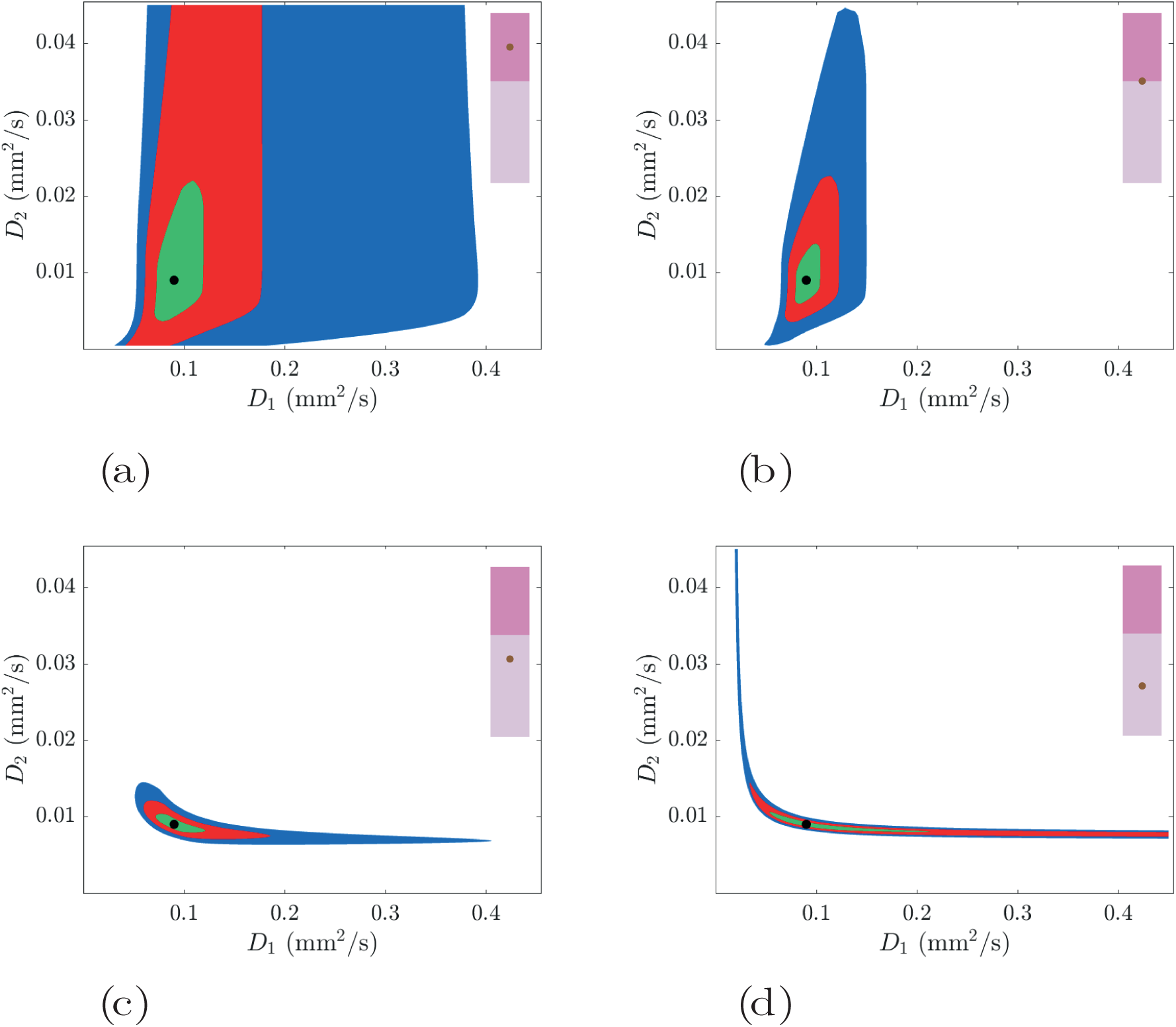
The role of probe location, *p*. As in Figure 3 the target parameter pair, 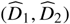, is highlighted with a black circle and the insets show various experimental designs with the brown circles showing the probe location relative to the tissue geometry. In each subfigure, the coloured regions satisfy *ℐ*_1_(*D*_1_, *D*_2_|*p*, *ε*) = 1 for *ε* = 0.5 (green), 1.0 (red) and 1.5 (blue). (**a**) Probe at the centre of the skin layer, *p* = *l*_1_/2; (**b**) Probe at the layer interface, *p* = *l*_1_; (**c**) Probe at the centre of the two-layer system, *p* = *l*_2_/2; (**d**) Probe at the centre of the fat layer, *p* = *l*_1_ + (*l*_2_ - *l*_1_)/2. In all cases we set *l*_1_ = 1.6 mm and *l*_2_ = 4 mm.

All discussion of the results in Figures 3-4 are so far based on qualitative visual interpretations of the extent of the coloured regions in these plots. To provide more quantitative insight we introduce a metric

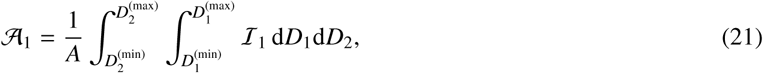

where *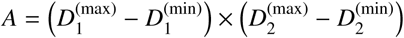* is the total area of the bounded parameter space in Figures 3-4. Here, 𝒜_1_ is the proportion of the bounded parameter space where the indicator function is unity when we consider data collected at a single probe. Although we write 𝒜_1_ in terms of a double integral in Equation (21), we find it simplest to interpret 𝒜_1_ as the proportion of the parameter space in which the indicator function is unity. Therefore, we estimate 𝒜_1_ by calculating *ℐ*_1_ at each point on the discretised (*D*_1_, *D*_2_) parameter space and computing the proportion of the 2001^2^ evaluations of *ℐ*_1_ that are unity. To interpret these results we note that smaller values of 𝒜_1_ are associated with improved experimental designs since the region of parameter space where *T* (*p*, *t*) is a close match to *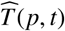* is reduced when 𝒜_1_ is smaller. Plotting 𝒜_1_ as a function of *p* in Figure 5 gives us greater quantitative insight into the role of probe placement.

**Figure 5:**
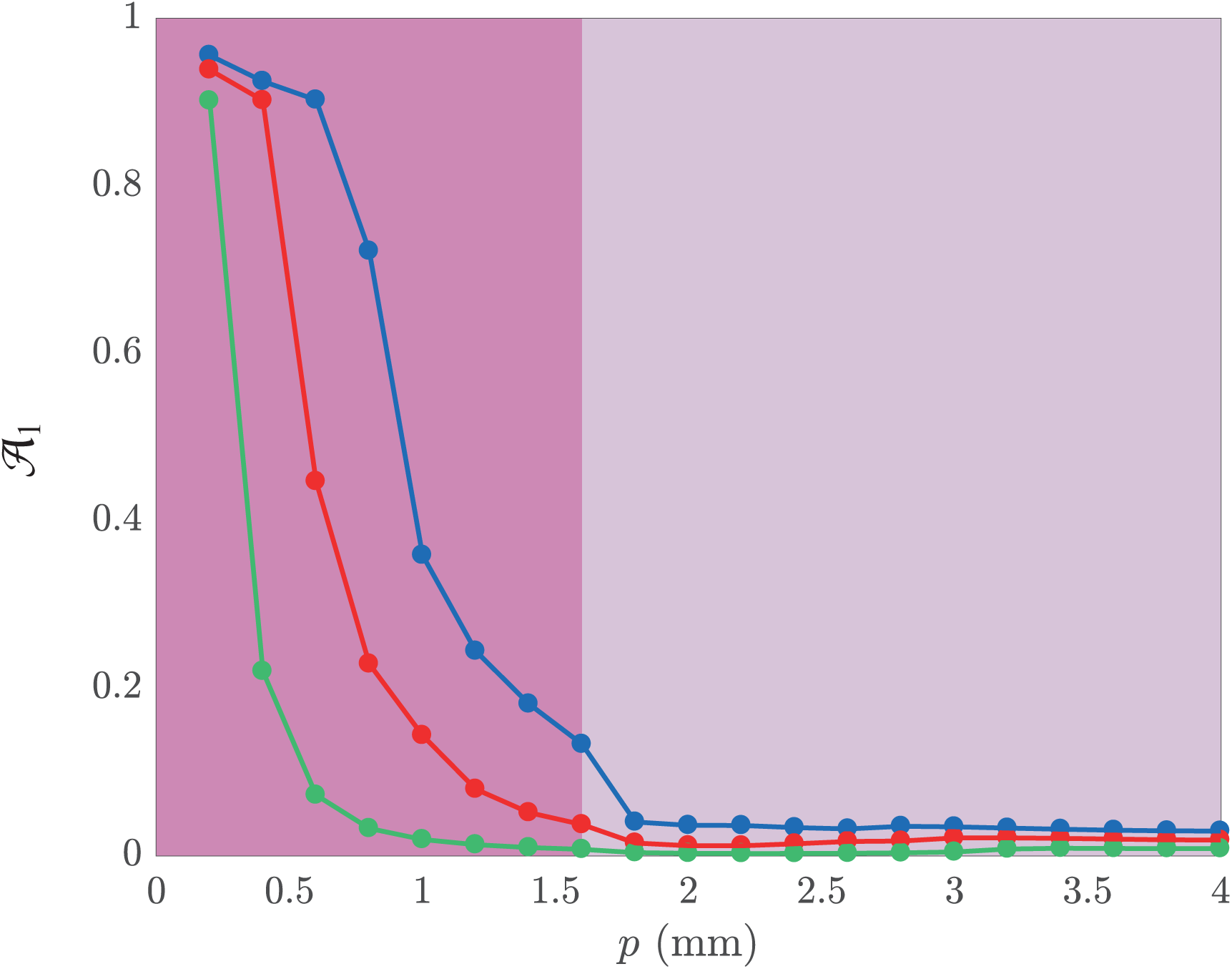
The influence of probe location, *p*, on 𝒜_1_ for *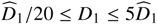* and *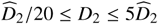*, and *l*_1_ = 1.6 mm and *l*_2_ = 4 mm. Plots of 𝒜_1_ are shown for *ε* = 0.5 (green), *ε* = 1.0 (red) and *ε* = 1.5 (blue). Calculations are performed for 20 equally-spaced values of *p*, from *p* = 0.2 mm to *p* = 4 mm. The bright pink and lighter pink background colours are chosen to correspond with the colour of the skin and fat layers in Figure 1(b).

Results in Figure 5 show that 𝒜_1_ appears to decrease with *p* for all values of *ε* we consider. The data in Figure 5 is useful because it provides a quantitative framework for examining the importance of the choice of probe placement, *p*. Overall we see that larger values of *p* lead to improved experimental designs, and we see that once *p* > 1.8 mm that 𝒜_1_ becomes relatively insensitive to any further increase in *p*. A simple recommendation we can provide from this exploration is that placing a single probe into the fat layer is a good experimental design.

All results presented in this study so far focus on the case where temperature data is recorded at a single location, *x* = *p*. While this constraint is an important feature of Cuttle’s experimental protocol [11–14], our mathematical modelling tools give us the flexibility to quantitatively explore the benefit of collecting data at more than one location in a controlled manner that is not possible experimentally. Therefore, we will now consider how our ability to estimate (*D*_1_, *D*_2_) are influenced if we were able to collect temperature data at two locations, *x* = *p* and *x* = *q*.

### 3.3. Parameter inference: two probes

To keep the presentation of our results manageable, when we consider the case where data is collected at two locations, *x* = *p* and *x* = *q*, we restrict our attention to the subset of cases where the location of the first probe, *x* = *p*, is fixed at *p* = *l*_2_ as in Cuttle’s experiments [11–14]. With this constraint, we then focus on how we might choose the location of the second probe, *x* = *q*. To achieve this we modify our definition of the indicator function to be

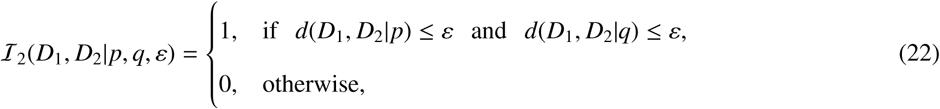

where *d*(*D*_1_, *D*_2_|*q*) is defined in exactly the same way as *d*(*D*_1_, *D*_2_|*p*) except that the spatial location is different. The key difference between *ℐ*_1_ and *ℐ*_2_ is that *ℐ*_2_ measures the closeness of *T* (*x*, *t*) and *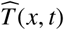* at both *x* = *p* and *x* = *q*, whereas *I*_1_ measures the closeness of *T* (*x*, *t*) and *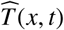* at *x* = *p* only. We follow our previous approach from Section by proposing four sensible choices for the placement of the second probe:

1. the centre of the skin layer, *q* = *l*_1_/2;

2. the layer interface, *q* = *l*_1_;

3. the centre of the two-layer system, *q* = *l*_2_/2; and

4. the centre of the fat layer, *q* = *l*_1_ + (*l*_2_ − *l*_1_)/2.

Results in Figure 6 show plots of the regions where *ℐ*_2_(*D*_1_, *D*_2_|*l*_2_, *q*, *ε*) = 1. The arrangement of the subfigures in Figure 6 corresponds to the arrangement of the subfigures in Figure 4 except that we now have two probes in the layered system. Comparing the extent of the coloured regions where *ℐ*_2_(*D*_1_, *D*_2_|*l*_2_, *q*, *ε*) = 1 in Figure 6 to the extent of the colored regions where *ℐ*_1_(*D*_1_, *D*_2_|*l*_2_, *ε*) = 1 in Figure 4 provides information about how the collection of additional data at a second location would improve our ability to reliably distinguish between *T* (*x*, *t*) and *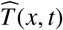* at *x* = *l*_2_ only (Figure 4), compared to our ability to distinguish between *T* (*x*, *t*) and *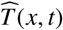* at both *x* = *l*_2_ and *x* = *q* (Figure 6). Overall, regardless of the choice of *q*, we see that working with a second probe always reduces the extent of the coloured region. Furthermore, comparing results across the four subfigures in Figure 6 indicates that the configuration in Figure 6(b), where the second probe is placed at the layer interface *x* = *l*_1_, is the best configuration of these four possibilities since the extent of the coloured regions is smallest.

**Figure 6:**
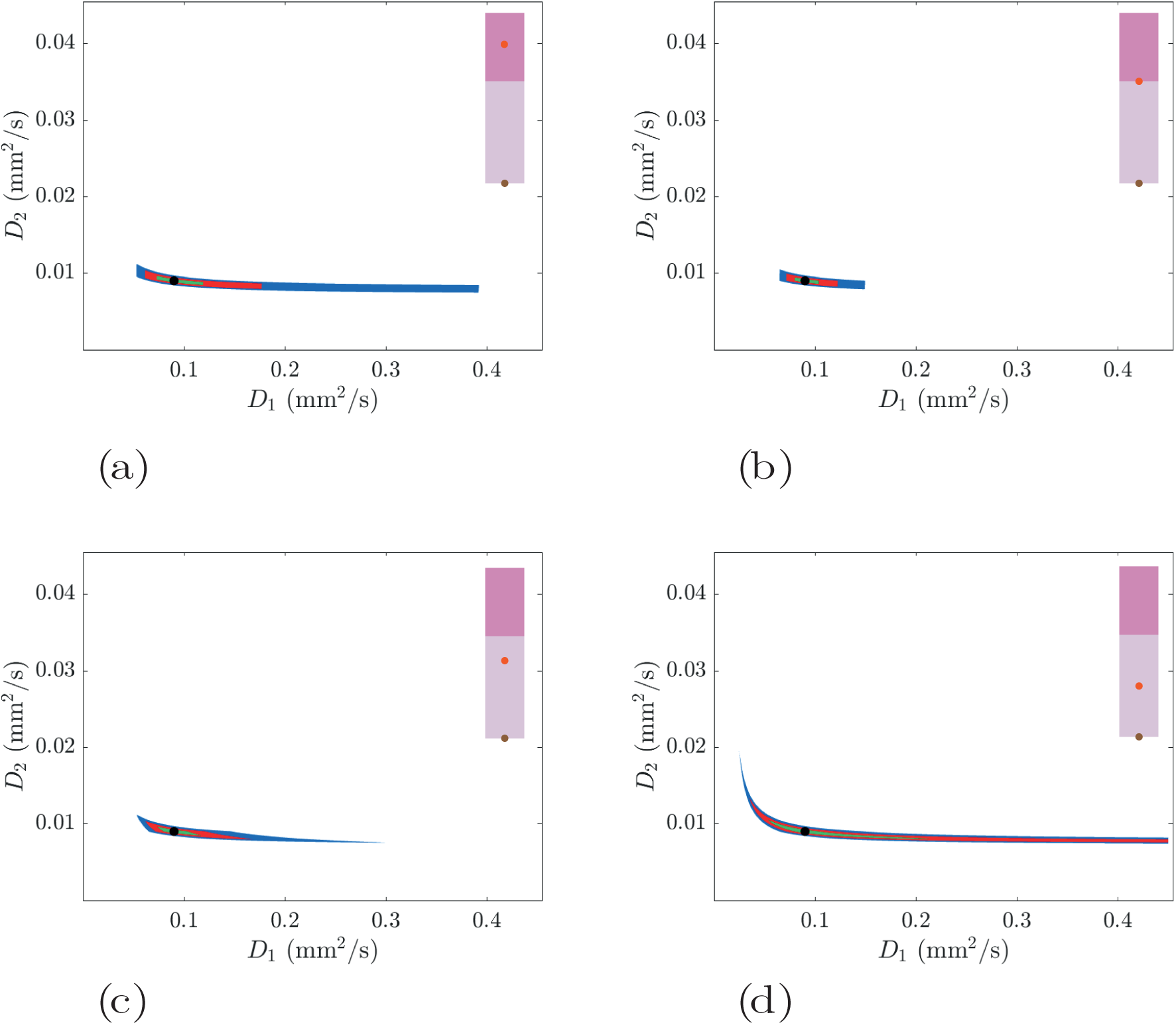
The role of the second probe location, *q*. As with Figure 4 the target parameter pair, 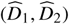, is highlighted with a black circle and the insets show various experimental designs with the brown circles showing the fixed first probe location and the red circles showing the variable second probe location relative to the tissue geometry. In each subfigure, the coloured regions satisfy *ℐ* _2_(*D*_1_, *D*_2_I*l*_2_, *q*, *ε*) = 1 for *ε* = 0.5 (green), 1.0 (red) and 1.5 (blue). (**a**) Second probe at the centre of the skin layer, *q* = *l*_1_/2; (**b**) Second probe at the layer interface, *q* = *l*_1_; (**c**) Second probe at the centre of the two-layer system, *q* = *l*_2_/2; (**d**) Second probe at the centre of the fat layer, *p* = *l*_1_ + (*l*_2_ - *l*_1_)/2. In all results we set *l*_1_ = 1.6 mm and *l*_2_ = 4 mm.

### 3.4. Parameter inference: optimal placement of second probe

To extend the results in Figure 6, we now explore whether there is some optimal placement of the second probe.

To explore this question we introduce

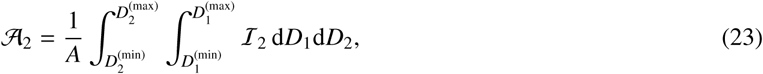

where 𝒜_2_ is the proportion of the parameter space that satisfies *ℐ*_2_(*D*_1_, *D*_2_|*l*_2_, *q*, *ε*) = 1. Similar to our approach in Section 3.2, we seek to find *q* which minimises 𝒜_2_. Results in Figure 7 show 𝒜_2_ as a function of *q*, for different choices of *ε*. Remarkably, we see that setting *q* = 1.6 mm minimises 𝒜_2_, for all *ε* considered. This results implies that the optimal location for a second probe, given that a first probe is already located at the bottom of the fat tissue *p* = *l*_2_, is at or near the layer interface, *q* = *l*_1_.

**Figure 7:**
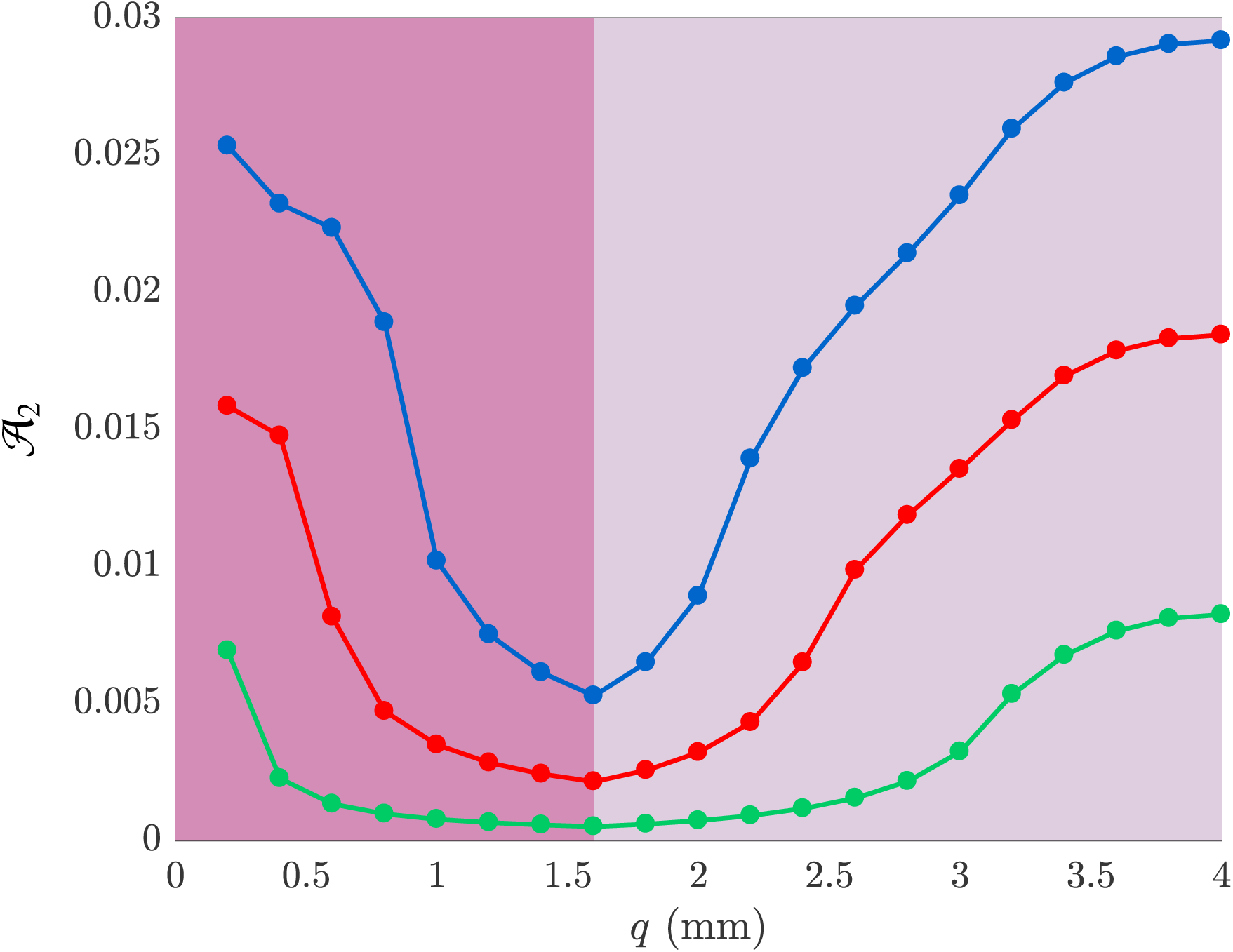
The influence of the second probe location, *q*, on 𝒜_2_ for *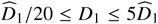* and s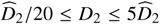, and *l*_1_ = 1.6 mm and *l*_2_ = 4 mm. Plots of 𝒜_2_ are shown for *ε* = 0.5 (green), *ε* = 1.0 (red) and *ε* = 1.5 (blue). In all cases *p* = *l*_2_ and calculations are performed for 20 equally-spaced values of *q* from *q* = 0.2 mm to *q* = 4 mm. The bright pink and lighter pink background colours are chosen to correspond with the colour of the skin and fat layers in Figure 1(b).

## 4. Conclusions and future directions

In this work we consider an experiential protocol designed by Cuttle and co-workers [11–14] to quantify the conduction of heat in living porcine (pig) tissues. This unique experimental protocol is very important because many experimental studies that examine heat conduction in skin tissue focus on non-living excised tissues [6, 7, 9, 10] whereas the protocol developed by Cuttle is far more realistic because they deal with living tissues, *in situ*. One of the constraints of Cuttle’s experimental protocol is that the temperature within the living tissues is monitored using a subdermal probe at a single location within the layered skin. Because skin is a layered structure, with the epidermis and dermis layers overlying a deeper fat layer, it is natural for us to model the conduction of heat in this system using a heterogeneous multilayer model where the thermal diffusivity in each layer can be different. In this work we idealise the skin tissues as a two-layer system with the upper layer representing the epidermis and dermis combined, and the lower layer representing the subdermal fat. Since one of the main biological functions of the fat layer is to provide thermal insulation [27], we expect that the thermal diffusivity of the fat layer to be different to the thermal diffusivity of the skin layer.

The key question we address in this work is to explore whether temperature data at a single location in a two-layer system is sufficient for us to reliably estimate the thermal diffusivity in the skin and fat layers, (*D*_1_, *D*_2_). Using biologically-motivated target values, 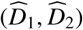, we solve the two-layer model and convert the spatiotemporal solution into a simple time series at a single location. This data is compatible with the kind of data recorded and reported by Cuttle and colleagues [11–14]. We then systematically scan the (*D*_1_, *D*_2_) parameter space, solving the model over four million parameter pairs, to explore the extent to which this time series data can be used to reliably identify the target parameters, 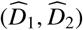. Our results show that our ability to estimate the parameters can be very sensitive using this kind of data as there are many combinations of parameter pairs, (*D*_1_, *D*_2_), leading to virtually indistinguishable time series data at a single location. Once we have demonstrated this sensitivity, we then explore the question of experimental design by using the mathematical model to explore the extent to which our ability to estimate (*D*_1_, *D*_2_) depends on the depth at which the subdermal probe is placed. In summary, we find that it is best to place the probe in the fat layer. This result is reassuring since Cuttle’s experimental protocol places the probe at the bottom of the fat layer [11–14]. We conclude by exploring the extent to which our ability to estimate (*D*_1_, *D*_2_) improves if we consider the case where two subdermal probes, placed at different locations, are used. Our results show that using a second probe always improves our ability to estimate (*D*_1_, *D*_2_), but there is still some sensitivity in terms of the placement of the subdermal probes. In summary, if it were possible to use two subdermal proves we find that given the first probe is placed at the bottom of the fat layer, and second probe ought to be placed at the interface of the skin and fat layers.

There are many ways that our study could be extended since we have invoked several simplifications and assumptions that could be relaxed. A key assumption in our work is that we treat the synthetic data generated by the mathematical model, *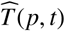* and 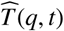, as being deterministic. This means that we neglect the role of experimental variability which is known to be important when dealing with biological data [39, 40]. If we had an estimate of the experimental variability in Cuttle’s measurements, we could incorporate this into our parameter sensitivity analysis by adding an appropriate noise signal, such as white noise, to *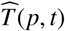* and 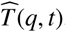, and then exploring how the incorporation of experimental variability influences our ability to estimate (*D*_1_, *D*_2_). Another feature of our mathematical model that could be explored further is our assumption that the boundary between the bottom of the fat layer and the underlying muscle and bone tissues, at *x* = *l*_2_ is perfectly insulating. In reality, we expect that there would be some transfer of heat from the fat tissues into the underlying muscle and bone, and this could be incorporated into the model using a Robin boundary condition. This approach would introduce an additional unknown heat transfer coefficient, thereby increasing the dimensionality of the parameter space to be explored. Both of these extensions could be considered in future studies.

Another natural extension of our current work would be to treat the conduction of heat in living skin as a three-layer problem instead of a two-layer problem. The three-layer problem could be constructed by treating the epidermis, dermis and fat as three distinct layers. While the semi-analytical solution strategy for solving the two-layer model generalises perfectly well to a three-layer model, the challenge of identifying three values of the thermal diffusivity instead of two values would become even more challenging when dealing with experimental observations where temperature is recorded at a single location in the layered system. Given that the main result of the current work highlights how challenging it can be to estimate parameters for a two-layer model, we anticipate that it is presently infeasible to meaningfully interpret data from Cuttle’s current experimental protocol using a three-layer model. However, if some of the current experimental constraints were to be alleviated and it became feasible to collect experimental temperature data at multiple positions simultaneously, then it is possible that working with a three-layer model could be reasonable in the future.

Overall, our work points to an important role for mathematical models and parameter estimation when dealing with complex biological processes that take place in complex biological environments. It is well-known that biology experiments can be difficult to reproduce and the issues associated with experimental reproducibility has important scientific and economic consequences [41]. Our view is that one way to start dealing with issues of experimental reproductivity in the biological sciences is to routinely apply mathematical models to quantitatively mimic experimental data and to explore questions of optimal experimental design [42, 43]. Working with mathematical models can be more transparent than complicated experimental protocols, and this is one of the reasons why all code and algorithms associated with this work are freely available at GitHub.

## 5. Acknowledgements

We thank Dr Leila Cuttle for introducing us to this biologically-motivated heat transfer problem, and we acknowledge the support from the Australian Research Council (DE150101137, DP170100474). We also thank the editor-in-chief for their assistance.

## References

[1] RL Sheridan. Burns a practical approach to immediate treatment and long-term care. Manson Publishing, London (2012).

[2] CJ Andrews, L Cuttle, MJ Simpson. Quantifying the role of burn temperature, burn duration and skin thickness in an *in vivo* animal skin model of heat conduction. Int J Heat Mass Transfer. 101 (2016) 542.

[3] A Abdullahi, S Amini-Nik, M Jeschke. Animal models in burn research. Cell Mol Life Sci. 71 (2014) 3241.

[4] W Meyer, R Schwarz, K Neurand. The skin of domestic mammals as a model for the human skin, with special reference to the domestic pig. Curr Probl Dermatol. 7 (1978) 39.

[5] W Montagna, JS Yun. The skin of the domestic pig. J Invest Dermatol. 42 (1964) 11.

[6] FC Henriques, AR Moritz. Studies of thermal injury: I. The conduction of heat to and through skin and the temperatures attained therein. A theoretical and an experimental investigation. Am J Pathol. 23 (1947) 530.

[7] AR Moritz, FC Henriques. Studies of thermal injury: II. The relative importance of time and surface temperature in the causation of cutaneous burns. Am J Pathol. 23 (1947) 695.

[8] TP Sullivan, WH Eaglstein, SC Davis, P Mertz. The pig as a model for human wound healing. Wound Repair Regen. 9 (2001) 6676.

[9] MA El-Brawany, DK Nassiri, G Terhaar, A Shaw, I Rivens, K Lozhken. Measurement of thermal and ultrasonic properties of some biological tissues. J Med Eng Technol. 33 (2009) 249.

[10] SM Brown, ML Baesso, J Shen, RD Snook. Thermal diffusivity of skin measured by two photothermal techniques. Anal Chim Acta 282 (1993) 711.

[11] L Cuttle, M Kempf, GE Phillips, J Mill, MT Hayes, JF Fraser, X-Q Wang, RM Kimble. A porcine deep dermal partial thickness burn model with hypertrophic scarring. Burns. 32 (2006) 806.

[12] L Cuttle, M Kempf, O Kravchuk, N George, P-Y Liu, H-E Chang, J Mill, X-Q Wang, RM Kimble. The efficacy of aloe vera, tea tree oil and saliva as first aid treatment for partial thickness burn injuries. Burns. 34 (2008) 1126.

[13] L Cuttle, M Kempf, O Kravchuk, GE Phillips, J Mill, X-Q Wang, RM Kimble. The optimal temperature of first aid treatment for partial thickness burn injuries. Wound Repair Regen. 16 (2008) 626.

[14] L Cuttle, M Kempf, P-Y Liu, O Kravchuk, RM Kimble.The optimal duration and delay of first aid treatment for deep partial thickness burn injuries. Burns. 36 (2010) 673.

[15] P Haridas, JA McGovern, DLS McElwain, MJ Simpson. Quantitative comparison of the spreading and invasion of radial growth phase and metastatic melanoma cells in a three-dimensional human skin equivalent model. PeerJ. 5 (2017) e3754.

[16] P Haridas, AP Browning, JA McGovern, DLS McElwain, MJ Simpson. Three-dimensional experiments and individual based simulations show that cell proliferation drives melanoma nest formation in human skin tissue. BMC Syst Biol. 12 (2018) 34.

[17] MJ Simpson, S McInerney, EJ Carr, L Cuttle. Quantifying the efficacy of first aid treatments for burn injuries using mathematical modelling and *in vivo* porcine experiments. Sci Rep. 7 (2017) 10925.

[18] HH Pennes. Analysis of tissue and arterial blood temperatures in the resting human forearm. J Appl Physiol. 1 (1948) 93.

[19] E Kengne, A Lakhssassi. Bioheat transfer problem for one-dimensional spherical biological tissues. Math Biosci. 269 (2015) 1.

[20] GN Mercer, HS Sidhu. A heat transfer model describing burns to the skin from automotive airbags. ANZIAM J. 47 (2006) 16

[21] KR Diller, LJ Hayes, GK Blake. Analysis of alternate models for simulating thermal burns. J Burn Care Rehab. 12 (1991) 177.

[22] A Baldwin, J Xu, D Attinger. How to cool a burn: a heat transfer point of view. J Burn Care Res 33 (2012) 176.

[23] DP Orgill, MG Solari, MS Barlow, NE OConnor. A finite-element model predicts thermal damage in cutaneous contact burns. J Burn Care Res. 19 (1998) 203.

[24] C Orndorff, S Ponomarev, w Dai, A Bejan. Thermal analysis in a triple-layered skin structure with embedded vasculature, tumor, and gold nanoshells. Int J Heat Mass Transfer. 111 (2017) 677.

[25] D Sarker, A Haji-Sheikh, A Jain. Temperature distribution in multi-layer skin tissue in presence of a tumor. Int J Heat Mass Transfer. 91 (2015) 602.

[26] MJ Simpson. Depth-averaging errors in reactive transport modeling. Water Res Resour. 45 (2009) W02505.

[27] MG Hayward,WR Keatinge. Roles of subcutaneous fat and thermoregulatory reflexes in determining ability to stabilize body temperature in water. J Physiol. 320 (1981) 229.

[28] EJ Carr, IW Turner, P Perré. Macroscale modelling of multilayer diffusion: Using volume averaging to correct the boundary conditions. Appl Math Model. 47 (2017) 600.

[29] EJ Carr, IW Turner. A semi-analytical solution for multilayer diffusion in a composite medium consisting of a large number of layers. Appl Math Model. 40 (2016) 7034.

[30] MR Rodrigo, A Worthy. Solution of multilayer diffusion problems via the Laplace transform. J Math Anal Appl. 44 (2016) 475.

[31] N Sheils, B Deconinck. Heat conduction on the ring: Interface problems with periodic boundary conditions. Appl Math Lett. 37 (2014) 107.

[32] N Sheils. Multilayer diffusion in a composite medium with imperfect contact. Appl Math Model. 46 (2017) 450.

[33] L Debnath, D Bhatta. Integral transforms and their applications. Chapman and Hall, Boca Raton (2007).

[34] KA Landman, MA McGuinness. Mean action time for diffusive processes. J Appl Math Dec Sci. 4 (2000) 125.

[35] EJ Carr, MJ Simpson. Accurate and efficient calculation of response times for groundwater flow. J Hydrol. 558 (2018) 470.

[36] MJ Simpson. Critical time scales for morphogen gradient formation: Concentration or gradient criteria? Int J Heat Mass Transfer. 106 (2017) 570.

[37] RI Hickson, SI Barry, GN Mercer. Critical times in multilayer diffusion. Part 1: Exact solutions. Int J Heat Mass Transfer. 52 (2009) 5776.

[38] RI Hickson, SI Barry, GN Mercer. Critical times in multilayer diffusion. Part 2: Approximate solutions. Int J Heat Mass Transfer. 52 (2009) 5784.

[39] w Jin, ET Shah, CJ Penington, SW McCue, PK Maini, MJ Simpson. Logistic proliferation of cells in scratch assays is delayed. Bull Math Biol. 79 (2017) 1028.

[40] DJ Warne, RE Baker, MJ Simpson. Optimal quantification of contact inhibition in cell populations. Biophys J. 113 (2017) 1920.

[41] w Jin, ET Shah, CJ Penington, SW McCue, LK Chopin, MJ Simpson. Reproducibility of scratch assays is affected by the initial degree of confluence: experiments, modelling and model selection. J Theor Biol. 390 (2016) 136.

[42] ST Johnston, ET Shah, LK Chopin, DLS McElwain, MJ Simpson. Estimating cell diffusivity and cell proliferation rate by interpreting IncuCyte ZOOMTM assay data using the Fisher-Kolmogorov model. BMC Syst Biol. 9 (2015) 38.

[43] ST Johnston, JV Ross, BJ Binder, DLS McElwain, P Haridas, MJ Simpson. Quantifying the effect of experimental design choices for in vitro scratch assays. J Theor Biol. 400 (2016) 19.

